# Prey range and genome evolution of *Halobacteriovorax marinus* predatory bacteria from an estuary

**DOI:** 10.1101/180265

**Authors:** Brett G. Enos, Molly K. Anthony, Joseph A. DeGiorgis, Laura E. Williams

**Affiliations:** Department of Biology, Providence College, Providence, RI; Cellular Dynamics Program, Marine Biological Laboratory, Woods Hole, MA

**Keywords:** predation, horizontal gene transfer, host range, marine ecosystem, mobile genetic element

## Abstract

**Background:** *Halobacteriovorax* are saltwater-adapted predatory bacteria that attack Gram-negative bacteria and therefore may play an important role in shaping microbial communities. To understand the impact of *Halobacteriovorax* on ecosystems and develop them as biocontrol agents, it is important to characterize variation in predation phenotypes such as prey range and investigate the forces impacting *Halobacteriovorax* genome evolution across different phylogenetic distances.

**Results:** We isolated *H. marinus* BE01 from an estuary in Rhode Island using *Vibrio* from the same site as prey. Small, fast-moving attack phase BE01 cells attach to and invade prey cells, consistent with the intraperiplasmic predation strategy of *H. marinus* type strain SJ. BE01 is a prey generalist, forming plaques on *Vibrio* strains from the estuary as well as *Pseudomonas* from soil and *E. coli*. Genome analysis revealed that BE01 is very closely related to SJ, with extremely high conservation of gene order and amino acid sequences. Despite this similarity, we identified two regions of gene content difference that likely resulted from horizontal gene transfer. Analysis of modal codon usage frequencies supports the hypothesis that these regions were acquired from bacteria with different codon usage biases compared to *Halobacteriovorax*. In BE01, one of these regions includes genes associated with mobile genetic elements, such as a transposase not found in SJ and degraded remnants of an integrase occurring as a full-length gene in SJ. The corresponding region in SJ included unique mobile genetic element genes, such as a site-specific recombinase and bacteriophage-related genes not found in BE01. Acquired functions in BE01 include the *dnd* operon, which encodes a pathway for DNA modification that may protect DNA from nucleases, and a suite of genes involved in membrane synthesis and regulation of gene expression that was likely acquired from another *Halobacteriovorax* lineage.

**Conclusions:** Our results support previous observations that *Halobacteriovorax* prey on a broad range of Gram-negative bacteria. Genome analysis suggests strong selective pressure to maintain the genome in the *H. marinus* lineage represented by BE01 and SJ, although our results also provide further evidence that horizontal gene transfer plays an important role in genome evolution in predatory bacteria.

## Background

Predation is an important force shaping microbial communities, which include microbial species that prey on other microbes. Eukaryotic microbial predators have received the majority of attention; however, bacterial predators are found in a wide range of environments and attack bacteria and fungi [1]. Predatory bacteria such as *Bdellovibrio bacteriovorus* attack animal and plant pathogens, which makes them a potential biocontrol agent and an alternative to antibiotics [2, 3]. To further understand bacterial predation and inform development of predatory bacteria as biocontrol agents, it is important to characterize variation in predation phenotypes such as prey range and examine evolution of predatory bacteria lineages at different scales.

*Halobacteriovorax* is a genus of predatory bacteria belonging to the Deltaproteobacteria. Similar to *Bdellovibrio bacteriovorus*, which is also a member of Deltaproteobacteria, *Halobacteriovorax* exhibits a biphasic life cycle [4, 5]. In the attack phase, small, highly motile predatory bacteria cells search for prey bacteria and attach to the prey cell envelope. The predatory cell then invades the prey periplasm and re-shapes the prey cell envelope to form a bdelloplast. In the subsequent growth phase, the predatory cell residing in the periplasm secretes lytic enzymes into the prey cytoplasm. The enzymes digest prey cell contents, and the predatory cell uses the prey components to build its own macromolecules. After depleting the prey cell cytoplasm, the predatory cell divides into multiple progeny, which secrete lytic enzymes to lyse the bdelloplast and release themselves to enter the attack phase.

Because of the similarity in predatory life cycle between *Halobacteriovorax* and *Bdellovibrio bacteriovorus*, *Halobacteriovorax* species were originally classified within the genus *Bdellovibrio*. Analysis of 16S rRNA gene sequences led to an initial reclassification into the genus *Bacteriovorax* [6] and then a subsequent reclassification into the genus *Halobacteriovorax* within the family Halobacteriovoraceae [7]. *Halobacteriovorax* is adapted to saltwater environments and is distributed worldwide in oceans, estuaries and saltwater lakes [8]. Analysis of gene sequences from *Halobacteriovorax* of different saltwater environments revealed multiple phylogenetic clusters or operational taxonomic units [9]. The *H. marinus* type strain SJ belongs to cluster III and was isolated over 25 years ago off the coast of St. John’s Island in the Caribbean [5].

As a widespread, albeit seasonally fluctuating, member of saltwater ecosystems, *Halobacteriovorax* may play an important role in shaping microbial communities at these sites. One experiment compared the impact of naturally occurring *Halobacteriovorax* versus naturally occurring marine bacteriophage on mortality of *Vibrio parahaemolyticus* added to microcosms of surface water samples [10]. *Halobacteriovorax* appeared to cause a larger reduction in *V. parahaemolyticus* cell density than bacteriophage. Studies of other ecosystems, such as the coral microbiome, have also suggested that *Halobacteriovorax* may impact microbial community structure [11].

How *Halobacteriovorax* shapes saltwater microbial communities depends in part on which bacterial species are susceptible to predation by different *Halobacteriovorax* strains. Tests of *Halobacteriovorax* isolates from various saltwater environments indicate that, in general, this genus has a broad prey range [12, 13]. For example, predatory bacteria in saltwater aquarium and tidal pool samples attacked a phylogenetically diverse set of prey, including multiple species of *Vibrio*, *Pseudomonas* and *E. coli* [13]. Other studies show that within the genus, some *Halobacteriovorax* isolates may have a narrower prey range; for example, *Halobacteriovorax* isolated from seawater attacked multiple strains of *V. parahaemolyticus* but did not attack two other *Vibrio* species, *E. coli* or *Salmonella* serovar Typhimurium [14]. The prey species used to initially isolate *Halobacteriovorax* from water samples likely biases which predatory strains are recovered and therefore affects our understanding of variation in prey range phenotypes. This was shown when *Halobacteriovorax* with broader prey ranges were isolated from a tidal river using *E. coli* or *Salmonella* serovar Typhimurium [14].

To understand the adaptation and evolution of *Halobacteriovorax*, it is important to examine genome evolution across a range of phylogenetic distances. Currently, *H. marinus* SJ is the only complete genome for family Halobacteriovoraceae [5]. Draft genomes are available for four strains representing four other phylogenetic clusters of *Halobacteriovorax* [15]. Overall, genes in *Halobacteriovorax* show high sequence divergence, illustrated by the large proportion of predicted genes with no significant matches to other genera [5] and a relatively low average amino acid identity among the five *Halobacteriovorax* genomes [15]. Genome evolution in *Halobacteriovorax* may be affected by horizontal gene transfer, with multiple regions of the *H. marinus* SJ genome showing signatures associated with foreign DNA [5]. The extent of horizontal gene transfer and its impact on functional capacity is unknown.

To further investigate phenotypic and genotypic variation in *Halobacteriovorax*, we isolated a strain of *H. marinus* from an estuary using a *Vibrio* strain from the same site. We tested the prey range of the isolate against bacteria from the estuary and bacteria from other environments to explore variation in this predation phenotype. Comparative genomics with the closely related type strain *H. marinus* SJ revealed two regions of gene content difference that likely arose via horizontal gene transfer.

## Results

### Small, fast-moving Halobacteriovorax marinus BE01 cells invade prey cells

We isolated a strain of predatory bacteria from an estuary in Rhode Island using a *Vibrio* strain from the same estuary as prey. The predatory bacteria isolate has two copies of the 16S rRNA gene, one of which is identical to that of *Halobacteriovorax marinus* type strain SJ, whereas the other copy differs at only one nucleotide position. This supports classification of the isolate as *H. marinus*, and we further distinguish it as strain BE01. *H. marinus* BE01 and *H. marinus* SJ have very similar cell morphologies. BE01 attack phase cells are small and highly motile (Figure 1a and Additional File 1). They have a characteristic vibroid (comma-shaped) morphology with a single polar flagellum (Figure 1b). *H. marinus* BE01 forms completely clear, uniform plaques on lawns of susceptible prey bacteria (Figure 1c). Observations by 1000x phase-contrast microscopy show that BE01 invades prey cells. The closely related type strain SJ occupies the periplasmic space of Gram-negative prey cells after invasion [5], suggesting that BE01 is also an intraperiplasmic predator.

**Figure 1.**
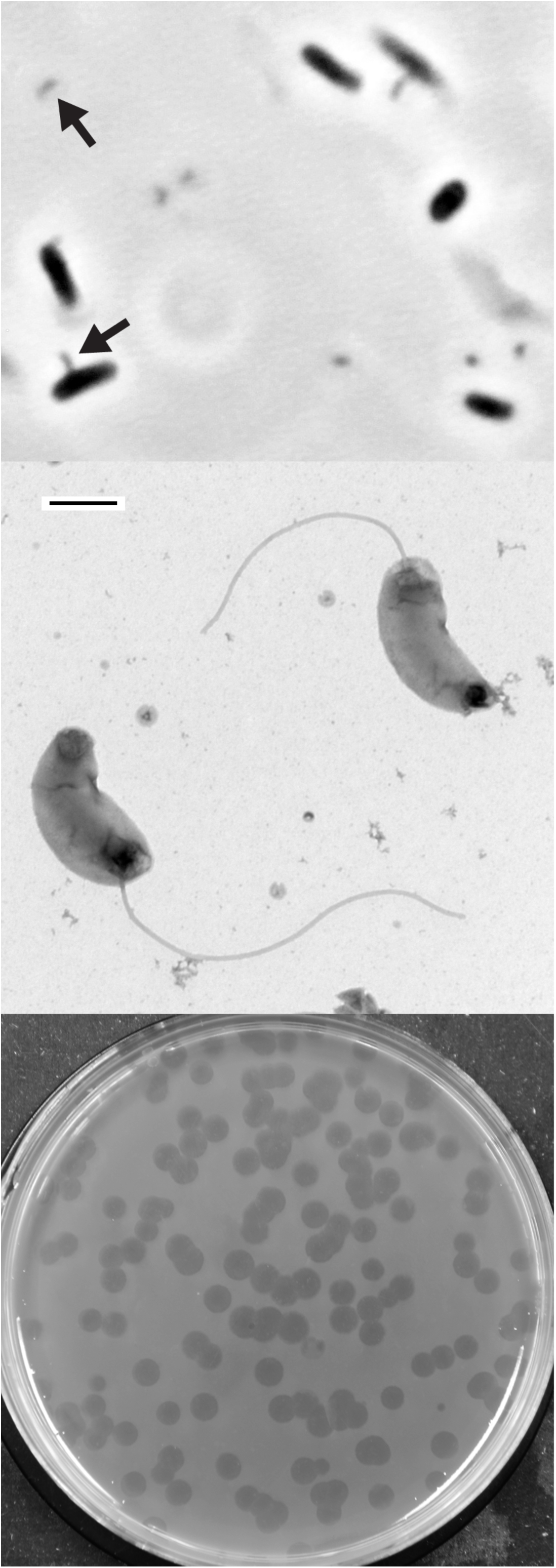
*Halobacteriovorax marinus* BE01 microscopy and plaque formation. (a) 1000x phase-contrast microscopy of small, comma-shaped BE01 cells (arrows) and larger *E. coli* ML35 cells. (b) BE01 cells stained with uranyl acetate and imaged with electron microscopy. Scale bar is 500 nm. (c) Plaques formed by BE01 on a lawn of *Vibrio* using double agar overlay.

### Halobacteriovorax marinus BE01 is a prey generalist

To assess prey range, we challenged *H. marinus* BE01 with different Gram-negative prey bacteria (Additional File 2). To test BE01’s ability to attack bacteria that it is likely to encounter in its natural habitat, we isolated multiple *Vibrio* strains from the estuary site and chose four distinct strains based on 16S rRNA gene sequences (Additional File 3). We also tested whether BE01 could attack Gram-negative isolates from other environments by challenging it with an *Acinetobacter* strain isolated from a freshwater lake, a *Pseudomonas* strain isolated from soil and two strains of *E. coli*, including ML35, a commonly used prey strain in studies of *Bdellovibrio*. We considered BE01 able to attack a particular prey strain if plaques formed on a lawn of that strain in a double agar overlay assay. Based on the results presented in Table 1, *H. marinus* BE01 appears to be a prey generalist, attacking all four *Vibrio* as well as *Pseudomonas* and both strains of *E. coli*. Plaque formation was consistent over three biological replicates.

**Table 1.** Prey range of *Halobacteriovorax marinus* BE01.

| Genus | Strain ID | Environment | Plaque formation? |
| --- | --- | --- | --- |
| *Vibrio* | 0024 | Estuary | Yes |
| *Vibrio* | 0026 | Estuary | Yes |
| *Vibrio* | 0027 | Estuary | Yes |
| *Vibrio* | 0028 | Estuary | Yes |
| *Acinetobacter* | 0036 | Freshwater | No |
| *Pseudomonas* | 0042 | Soil | Yes |
| *Escherichia* | 0057 |  | Yes |
| *Escherichia* | ML35 |  | Yes |

### Halobacteriovorax marinus BE01 genome is highly similar to SJ, but lacks plasmid

Table 2 shows general statistics for the chromosomes of *H. marinus* BE01 (CP017414) and SJ (NC_016620). The chromosome sequences of these two strains are very similar in size and identical in GC content. Average nucleotide identity (ANI) between the two strains is 98.2% when calculated by JSpecies using nucmer [16] and 98.0% when calculated at http://enve-omics.ce.gatech.edu/ani/ [17]. Initially, we annotated the BE01 chromosome using the Prokaryotic Genome Annotation Pipeline (PGAP) at GenBank and compared it to the existing GenBank annotation of SJ. PGAP classified more protein-coding genes as hypothetical proteins in BE01 compared to SJ (2,398 versus 1,571). Some of these classifications in the BE01 chromosome appear overly conservative; for example, BIY24_00015 in strain BE01 is annotated as a hypothetical protein although the amino acid sequence is 99% identical to BMS_0003 in strain SJ, which is annotated as DNA recombination protein RecF on the basis of conserved protein domain families. We therefore submitted both BE01 and SJ chromosome sequences to the Rapid Annotation using Subsystem Technology (RAST) server for annotation [18–20]. RAST classified a similar number of protein-coding genes as hypothetical proteins in the two strains (Table 2), and the proportion of hypothetical proteins was closer to the PGAP annotation of SJ. We supplemented the RAST annotation with Infernal annotation [21] to detect RNA-coding sequences and proceeded with our analyses using the RAST+Infernal annotations, which can be found as text files at the figshare repository (https://figshare.com/projects/Supporting_data_for_Halobacteriovorax_BE01_paper/242 29).

**Table 2.**
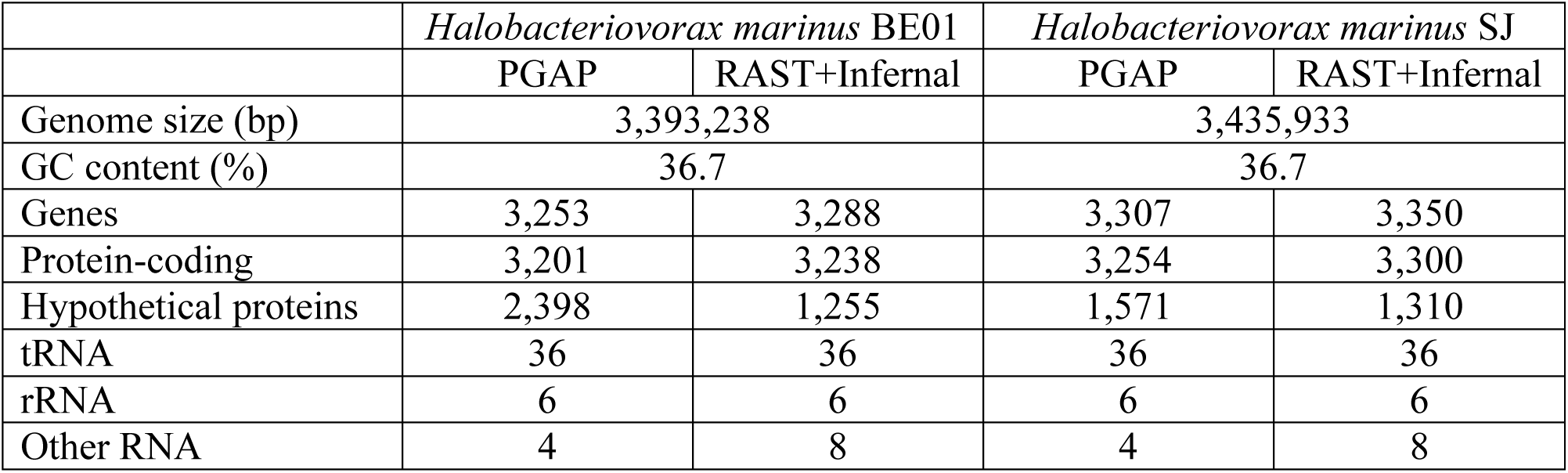
Chromosome statistics.

The genome of *H. marinus* SJ includes a small (1,973 bp) plasmid with a single coding sequence [5]. To determine whether *H. marinus* BE01 harbors a plasmid, we used megablast to identify the top hits for each of the 93 contigs generated by *de novo* assembly using PacBio reads. With the exception of the contig corresponding to the BE01 chromosome, all contigs aligned with at least 97% similarity to *E. coli* sequences in the non-redundant GenBank database. This is expected because we did not separate predatory bacteria cells from *E. coli* prey cells before extracting genomic DNA. Based on the megablast results, we conclude that the *H. marinus* BE01 genome consists of one chromosome and no plasmids.

### Conservation of synteny and amino acid sequences between H. marinus genomes

Using RAST, we identified 3,048 bidirectional best hits between BE01 and SJ. To check the accuracy of the RAST analysis, we used the Reciprocal Smallest Distance algorithm [22], which detected 3,040 orthologs. By plotting the position of the RAST bidirectional best hits on each chromosome, we observed extremely high conservation of gene order between the two *H. marinus* strains (Figure 2a). We did not detect any major inversions, translocations or duplications. Most bidirectional best hits between BE01 and SJ have high amino acid sequence identity. In particular, 86% of bidirectional best hits (2,610/3,048) have at least 98% amino acid identity, and 94% (2,865/3,048) have at least 96% amino acid identity (Figure 3). Only 27 bidirectional best hits have <70% identity at the amino acid sequence level, and many of these genes occur in one of the two major regions of difference detected in the synteny plot (Figure 2b). Such high conservation of gene order and amino acid sequence across the chromosome suggests that the lineage of *Halobacteriovorax marinus* represented by these two strains is experiencing strong purifying selection.

**Figure 2.**
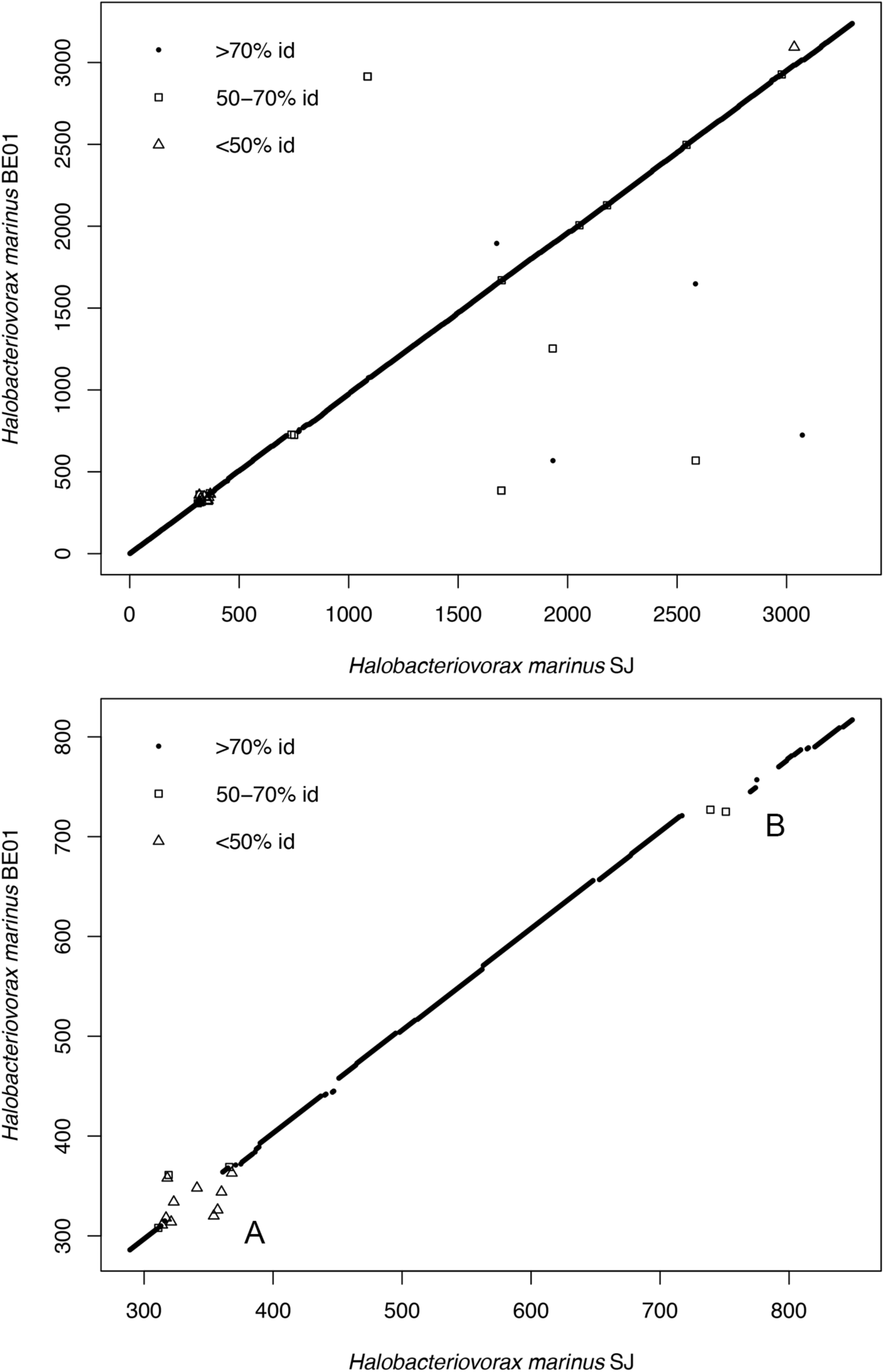
Synteny plot of bidirectional best hits between *H. marinus* BE01 and SJ. Bidirectional best hits identified by RAST are plotted based on their gene number on each chromosome. Individual genes are denoted with symbols corresponding to the similarity between BE01 and SJ amino acid sequences. (a) shows the entire chromosomes, whereas (b) highlights the two major regions of difference in gene content (labeled A and B).

**Figure 3.**
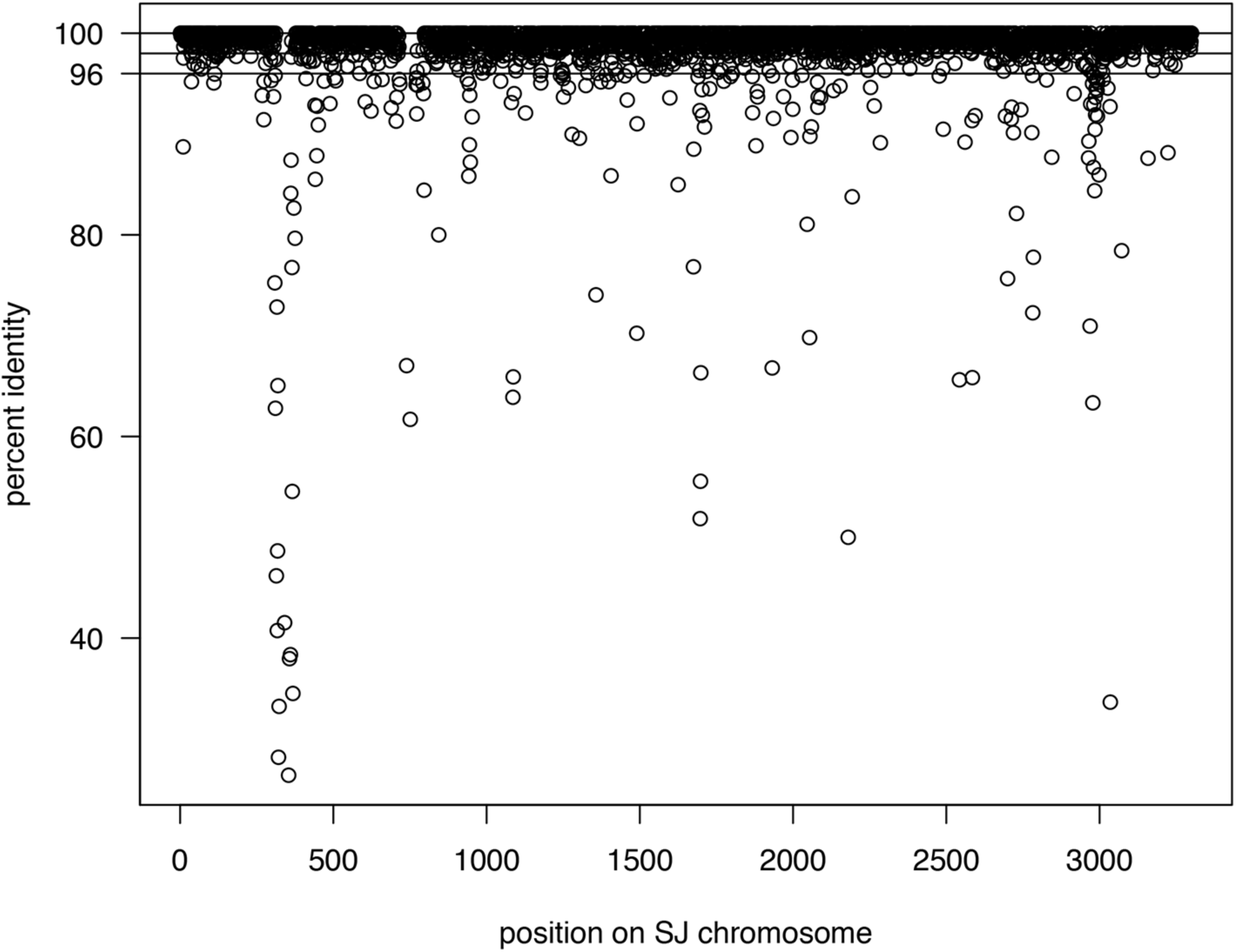
Amino acid identity of bidirectional best hits. Bidirectional best hits identified by RAST are plotted based on their position on the SJ chromosome and the similarity between BE01 and SJ amino acid sequences. Horizontal lines indicate 100%, 98% and 96% amino acid identity.

### Differences in gene content between H. marinus genomes

The synteny plot of bidirectional best hits revealed two major regions of difference in gene content between *H. marinus* BE01 and *H. marinus* SJ (Figure 2). One of these regions (region B in Figure 2b) is bounded by a hypothetical protein (BE01_721 and SJ_717, see Additional File 4 for corresponding PGAP locus tags) and a TonB-dependent outer membrane receptor (BE01_770 and SJ_792). In BE01, region B encompasses 48 genes, 32 of which (67%) are unique to BE01, whereas in SJ, this region encompasses 74 genes, 56 of which (76%) are unique to SJ. In BE01, seven of the 48 genes are unidirectional best hits against the SJ genome, with <65% amino acid identity, and nine of the 48 genes are bidirectional best hits, with >60% amino acid identity.

Regarding functions annotated in region B in the BE01 genome, 23 of the 48 genes (48%) are hypothetical proteins with no predicted function. Three genes are annotated with functions related to horizontal gene transfer. BE01_722 is annotated as a mobile element protein, with hits to a COG (COG3464) and a PFAM domain (pfam01610) for a transposase family. Two consecutive genes BE01_727-728 are both annotated as integrases. BE01_727 is a bidirectional best hit to SJ_739, whereas BE01_728 is a unidirectional best hit for the same SJ gene. BLASTX analysis of the nucleotide sequence spanning these two genes and the intergenic regions suggests that the two genes are pseudogenes of the full-length integrase. Accumulation of mutations has degraded the gene, leaving two frameshifted ORFs that align to different regions of the full-length SJ integrase sequence with 67% and 57% amino acid identity by BLASTP.

The presence of genes associated with mobile genetic elements led us to examine the genes unique to BE01 in this region, which may be the result of horizontal gene transfer events. We found two sets of genes indicative of HGT. One set of five genes (BE01_733-737) encodes *dnd* genes involved in phosphorothioation of DNA. The *dnd* operon is not found in *H. marinus* SJ, but it is found in multiple divergent bacterial lineages, with phylogenetic evidence suggesting horizontal transfer [23, 24]. We attempted to identify a likely source of the BE01 *dnd* operon, but each Dnd protein aligned with <55% identity to protein sequences in the database and had a different bacterial species as the top hit in BLASTP analysis, thereby providing no clear evidence of the donor species.

In addition to the *dnd* operon, we identified a set of nine genes (BE01_761-769) that may have been acquired from another *Halobacteriovorax* lineage. By BLASTP analysis, each of the amino acid sequences has 37-64% identity (query coverage ≥97%) to sequences in *Halobacteriovorax* sp. BAL6_X, which belongs to a different phylogenetic cluster than SJ and BE01. The nine genes are in the same order and orientation in BE01 and BAL6_X and include three genes involved in fatty acid and phospholipid metabolism and two genes encoding proteins with similarity to RNA polymerase sigma factor RpoE and an anti-sigma factor.

We also examined genes unique to SJ in region B to identify possible HGT events experienced by this strain. Forty-six of the 56 unique SJ genes were annotated as hypothetical proteins, with no predicted function. The remaining ten genes included six genes associated with mobile genetic elements, including a site-specific recombinase (SJ_729), an RNA-directed DNA polymerase (SJ_745) and four consecutive genes encoding a phage transcriptional regulator (SJ_755) and a Type I restriction-modification system (SJ_756-758). Overall, analysis of region B in BE01 and SJ suggests that it may be a “hotspot” for incorporation of mobile genetic elements in this lineage of *H. marinus*.

The other major region of difference (region A in Figure 2b) is bounded by chaperone protein DnaK (BE01_310 and SJ_313) on one end. On the other end, this region is bounded by different mannosyltransferases (BE01_364 or SJ_375), which are not each other’s bidirectional best hit. In BE01, region A encompasses 53 genes, 29 of which (55%) are unique to BE01, whereas in SJ, this region encompasses 61 genes, 25 of which (41%) are unique to SJ. In BE01, 12 of the 53 genes are unidirectional best hits against the SJ genome, with ≤40% amino acid identity, and 12 are bidirectional best hits, only two of which have amino acid identity >50%. In contrast to region B, which contains mostly unique gene content in each *H. marinus* strain, region A appears to have experienced more recombination and divergence of shared gene content (Figure 2b).

Regarding functions annotated in region A in the BE01 genome, 15 of the 53 genes (28%) are hypothetical proteins, with no predicted function. Among the remaining genes, we identified 22 genes with transferase activity, either annotated as transferases by RAST or classified as a transferase by analysis with InterProScan (GO term or detailed domain signature match). Ten of these transferase genes are unique to BE01. We also identified six genes involved in polysaccharide biosynthesis, three of which are unique to BE01.

We did not detect genes associated with mobile genetic elements, such as transposases or integrases, in this region in BE01.

Overall, these two regions encompass 61 of the 147 total unique genes (41%) in BE01 and 81 of the 187 total unique genes (43%) in SJ. A large proportion of unique genes across the whole chromosome are annotated as hypothetical proteins (70% in BE01 and 73% in SJ). These ORFs may not encode functional proteins, therefore the number of unique protein-coding genes may be even lower. This emphasizes the high degree of shared gene content between *H. marinus* BE01 and *H. marinus* SJ, with the two regions described above encompassing the majority of unique or highly divergent genes.

### Modal codon usage indicates horizontal gene transfer in regions of difference

To further explore the possibility of horizontal gene transfer in this lineage of *H. marinus*, we analyzed modal codon usage frequencies in both BE01 and SJ. Codon usage bias, in which certain codons are preferred for a particular amino acid, differs among bacterial species. Within a bacterial chromosome, regions with a codon usage bias that differs from that of the rest of the chromosome may have been horizontally transferred, although this is not the only explanation [25]. Here, we analyzed modal codon usage, which describes the codon usage frequencies of the largest number of genes in a given sequence [26]. We compared the entire chromosomes of BE01 and SJ and found a very small distance between the modes of codon usage frequencies (Table 3). This is expected based on the high average nucleotide identity between these two strains.

**Table 3.** Modal codon usage of *H. marinus* chromosomes and regions of gene content difference within each chromosome.

| Comparison | Sequence 1 | Sequence 2 | Distance between Sequence Modes | Distance between Modes of Shuffled Sequences |
| --- | --- | --- | --- | --- |
| Whole chromosomes | BE01 | SJ | 0.0241 | 0.0402 ± 0.0049 |
| BE01 regions | Region A | Chromosome (excluding regions A and B) | 0.3589 | 0.1047 ± 0.0117 |
| | Region B | Chromosome (excluding regions A and B) | 0.2976 | 0.1149 ± 0.0171 |
| | Region A | Region B | 0.2607 | 0.1429 ± 0.0257 |
| SJ regions | Region A | Chromosome (excluding regions A and B) | 0.3756 | 0.0985 ± 0.0092 |
| | Region B | Chromosome (excluding regions A and B) | 0.2237 | 0.0913 ± 0.0133 |
| | Region A | Region B | 0.3053 | 0.1385 ± 0.0166 |

To test our hypothesis that the regions of gene content difference discussed above were acquired by horizontal gene transfer from a bacterial species with a different codon usage bias, we performed within-genome pairwise comparisons to determine the distances between the modes of codon usage frequencies for the two regions and the rest of the chromosome (excluding these two regions). Each pairwise comparison included a test of the null hypothesis that the two sequences shared the same modal codon usage frequencies. This is done within the software package by combining all the genes from both sequences in a pairwise comparison and then generating two new random sets of genes with the same number of genes as the original. The software then calculates the modal codon usage frequencies of ten of these “shuffled” sequence pairs, generating an average and standard deviation. The average distance of these ten shuffled sequence pairs can be considered the expected distance if the two sequences shared the same modal codon usage frequencies [26].

Using this approach, we found that distances between the modes of codon usage frequencies for each region and the rest of the chromosome were larger than expected in both BE01 and SJ (Table 3). This supports the hypothesis that these regions were acquired via horizontal gene transfer from bacterial species with different codon usage biases compared to *H. marinus*. Highly expressed genes may also have different codon usage biases; however, given the annotated functions of genes in these regions, it is unlikely that this explains the distances observed. We also tested the distance between the modes of codon usage frequencies for the two regions themselves. These distances were also larger than expected in both BE01 and SJ (Table 3). This suggests that these regions were not acquired from a single bacterial species.

## Discussion

Based on prey range tests, *H. marinus* BE01 appears to be a prey generalist. It is capable of attacking *Vibrio* species isolated from the same estuary site as well as *Pseudomonas* from soil and two strains of *E. coli*, including ML35. We have used the same strain of *Pseudomonas* to isolate *Bdellovibrio* from soil (unpublished observations), and *E. coli* ML35 is often used to culture *Bdellovibrio* isolated from both freshwater and soil (for example, [27]). Our finding that *H. marinus* BE01 can attack these strains contrasts with reported observations that saltwater-adapted predatory bacteria generally do not attack the same prey species as *Bdellovibrio* [5]. Characterizing variation in predation phenotypes such as prey range is important for understanding diversity and adaptation in predatory bacteria. In this case, we used a *Vibrio* strain from the same site to isolate *H. marinus* BE01 rather than known strains such as *V. parahaemolyticus* P-5. This is a useful strategy for increasing the diversity of predatory bacteria recovered from different environments.

Comparison of the genomes of *H. marinus* BE01 and SJ clearly demonstrate that these strains are very closely related. The high conservation of synteny, gene content and nucleotide sequence is striking given the geographic distance between the sampling sites and the amount of time between sample collections. In particular, the lack of genome rearrangements and the large proportion of amino acid sequences with at least 96% identity between the two strains suggest strong selective pressure to maintain the genome within this lineage of *H. marinus*.

Despite this selective pressure, there are two genomic regions with evidence of past horizontal gene transfer events. Comparison of modal codon usage frequencies within these regions and the rest of the *H. marinus* chromosomes supports a hypothesis that these regions were acquired from bacterial species with different codon usage biases compared to *Halobacteriovorax*. One of the regions (region B) has a high proportion of unique gene content in both *H. marinus* BE01 and *H. marinus* SJ, including genes associated with mobile genetic elements such as transposons and bacteriophage.

Within region B, one set of nine genes was likely acquired from another *Halobacteriovorax* lineage. The donor may belong to *Halobacteriovorax* phylogenetic cluster X, since the top BLASTP hit for each of the BE01 genes is the sequenced representative of this cluster [15]. However, the current database is limited to only five *Halobacteriovorax* genomes. Without more genome data, it is unclear whether this suite of genes was exchanged directly between BAL6_X and BE01 and then diverged (thereby explaining <65% amino acid identity), or if it has been exchanged widely among other *Halobacteriovorax*, accumulating mutations in each new host. In the latter scenario, sequencing of additional *Halobacteriovorax* isolates from multiple phylogenetic clusters may identify a lineage with genes more similar in sequence to those of BE01. It is also unclear how these genes affect BE01’s functional capacity. Based on their annotations, they may affect membrane synthesis and regulation of gene expression.

Region B in BE01 also includes the *dnd* operon, which encodes a pathway for DNA modification. The *dnd* operon is found in multiple bacterial lineages. Phylogenies of individual *dnd* genes and investigations of genomic context suggest acquisition via horizontal gene transfer [23, 24]. Operons that include *dndA* typically have two orientations: *dndA* divergently transcribed from *dndBCDE* or *dnaAEDCB* transcribed in the same direction [28]. The operon in BE01 has the latter orientation, which may be less common based on genome surveys [28]. Although our analysis did not reveal a likely source of the *dnd* operon in *H. marinus* BE01, this operon has been reported in coastal Vibrionaceae and metagenome data of ocean samples [23]. The *dnd* genes in BE01 are intact, albeit highly divergent from *dnd* genes in the GenBank database. In other bacteria, Dnd proteins are involved in phosphorothioation, in which sulfur replaces a non-bridging oxygen molecule in the phosphate of the DNA backbone [29, 30]. Some researchers have suggested that phosphorothioation may protect genomic DNA from degradation by nucleases [29]. Predatory bacteria such as *Halobacteriovorax* rely on nucleases to digest prey DNA. If functional in BE01, the *dnd* operon may provide a horizontally acquired self-defense mechanism for *H. marinus* BE01 to protect its DNA from its own nucleases.

Similar to comparative genomics studies of *Bdellovibrio* [31–33], our analysis implicates an important role for horizontal gene transfer in the evolution of saltwater-adapted *Halobacteriovorax*. The extent to which predatory bacteria acquire genes from prey bacteria during predation is an interesting open question. Intraperiplasmic predators such as *Halobacteriovorax* and *Bdellovibrio bacteriovorus* secrete nucleases into the prey cytoplasm to digest genomic DNA; however, it is possible that partially digested fragments could be incorporated into the genome of the predatory bacteria cell during intraperiplasmic growth. It is also possible that partially digested fragments are released upon lysis of the prey cell by predatory progeny, enabling predatory bacteria cells in close proximity to take up the fragments and incorporate them into their chromosome.

In addition to unique genes suggestive of horizontal gene transfer, we also examined shared gene content between *H. marinus* BE01 and SJ. The high amino acid identity observed between these two strains is contrasted by the divergence of these sequences compared to database sequences, as described by Crossman and colleagues [5] in their analysis of the SJ genome. We also observed a high proportion of hypothetical proteins or proteins of unknown function. RAST annotations identified 39% of BE01 protein-coding genes and 40% of SJ protein-coding genes as hypothetical proteins. This emphasizes the need for characterization of these predicted genes to determine whether they encode a protein, and, if so, the function of that protein.

With a broad prey range such as observed here with *H. marinus* BE01, *Halobacteriovorax* species may exert a significant impact on microbial community structure in ecosystems such as estuaries. How these predatory bacteria affect nutrient cycling and fit into food web interactions is a key question for understanding these ecosystems [34]. In addition, predatory bacteria and *Halobacteriovorax* in particular have shown promise as an alternative to antibiotics in the control of bacterial pathogens [35]. Characterization of phenotypic and genotypic variation in a diverse range of *Halobacteriovorax* provides important information to advance development of these bacteria as biocontrol agents.

## Conclusions

The results of prey range tests of *H. marinus* BE01 are consistent with previous observations that *Halobacteriovorax* prey on a broad range of Gram-negative bacteria. Comparative genomics between *H. marinus* BE01 and the closely related *H. marinus* type strain SJ suggest strong selective pressure to maintain the genome in this *H. marinus* lineage. Despite this selective pressure, presence of mobile genetic element genes and atypical modal codon usage frequencies suggest that horizontal gene transfer impacted two genomic regions in this *Halobacteriovorax* lineage. HGT events provide these strains with unique functional capacities that may impact their adaptation.

## Methods

### Isolation and classification of environmental bacteria from estuary for use as prey

We isolated bacteria from Mount Hope Bay, an estuary in Bristol, RI (41.69717, - 71.24578) for use as potential prey. We collected water from 1 meter below the surface in sterile sample bottles and then filtered 100 ml through a 0.45 μm 47 mm membrane filter. We placed the filter in a 50 mm Petri dish on an absorbent pad presoaked with either 2 ml of sea water yeast extract (SWYE) broth [36] or Luria-Bertani (LB) broth (also known as lysogeny broth) with 3% NaCl. We incubated the filters at 29°C, then picked colonies and streaked them onto plates of the same growth medium used to presoak the filter. We performed four rounds of streak plates to ensure pure isolates. Using a similar approach, *Acinetobacter* #0036 was isolated from a freshwater lake, and *Pseudomonas* #0042 was isolated from soil.

To classify these isolates, we performed PCR targeting the 16S rRNA gene. We used primers 63F [37] and 1378R [38] with KAPA HiFi (high fidelity) DNA polymerase. PCR cycle conditions were: 95°C for 5 minutes followed by 30 cycles of 95°C for 30 seconds, 50°C for 30 seconds and 72°C for 2 minutes and a final extension step of 72°C for 10 minutes. After confirming presence of a PCR product by gel electrophoresis, we purified PCR products using the Ultra Clean PCR Clean-Up Kit (Mo Bio) and quantified them on a NanoDrop spectrophotometer. Sanger sequencing used the same primers as amplification and was performed by GeneWiz (South Plainfield, NJ). We used Phred/Phrap/Consed [39–41] to trim and assemble the reads, and we classified sequences using BLAST [42], the SILVA Incremental Aligner [43] and the Ribosomal Database Project classifier [44]. Additional File 2 shows the complete results of the classifications.

### Isolation of Halobacteriovorax marinus strain BE01

To isolate predatory bacteria, we collected a water sample (31 ppt salinity measured with a refractometer) from the same estuary site as described above following the same procedure. We combined 20 ml with 1 ml of *Vibrio* strain #0024 at 10^9^ cfu/ml and then incubated this enrichment at 26°C and 200 rpm. Enrichments were examined daily for 2 to 4 days by 1000x phase contrast microscopy for the presence of small, highly motile cells. Once we observed the presence of predatory bacteria, we filtered the enrichment through a 0.45 μm filter. We performed a 10-fold serial dilution of the filtrate in sterile 100% Instant Ocean (IO). Dilutions were plated using a double agar overlay method. Specifically, we added 1 ml of *Vibrio* strain #0024 at 10^9^ cfu/ml to test tubes containing 3.3 ml of molten Pp20 top agar (1 g polypeptone peptone and 19.5 g agar dissolved in 1 L of 70% IO). We vortexed to mix, then added 5 ml of the filtrate dilution to be plated and vortexed again. We poured this mixture onto Pp20 plates (1 g polypeptone peptone and 15 g agar dissolved in 1 L of 70% IO), allowed the top agar to solidify at room temperature and then incubated plates at 25°C. To check for possible bacteriophage, we examined the plates after 24 hours for plaques, but did not detect any. We observed plaques after 3-4 days. We picked plaques and made a lysate for each by placing a plaque in 20 ml of 100% IO with 1.5 ml of a *Vibrio* #0024 overnight culture. We incubated the lysates at 26°C and 200 rpm. After at least 24 hours of incubation, we used 1000x phase contrast microscopy to check for small, highly motile cells. After detecting predatory bacteria cells, we filtered the lysate through a 0.45 μm filter and repeated the double agar overlay technique to obtain individual plaques on a lawn of *Vibrio* #0024. The double agar overlay and plaque picking procedure was performed a total of three times to ensure a pure isolate of predatory bacteria. The lysate made from the final plaque picking procedure was filtered through a 0.45 μm filter. We combined 500 μl of this filtrate (containing cells of the pure isolate of predatory bacteria) with 500 μl sterile 50% glycerol and stored this stock at −80°C.

### Prey range tests

To obtain active *H. marinus* BE01 for prey range tests, we added a small amount of the −80°C stock to 15 ml of 100% IO mixed with 1 ml of an *E. coli* #0057 overnight culture. We chose to use *E. coli* because prior work reported viable *Vibrio* passing through 0.45 μm filters as minicells [14], which could confound the results of the prey range tests. We incubated the lysate at 26°C and 200 rpm. After three days, we filtered the lysate using a 0.45 μm filter to separate predatory bacteria from prey bacteria and cell debris. Swabs of the filtrate on LB plates confirmed that no viable *E. coli* passed through the filter. We performed 1:10 serial dilutions of the filtrate in 100% IO. To test prey range, we used the double agar overlay method described above to observe plaque formation. We cultured prey strains in 35 ml of SWYE broth for *Vibrio* prey strains or tryptic soy broth (TSB) for all other prey strains. We centrifuged cultures at 6000 rpm for 10 minutes, washed pellets in 100% IO and then resuspended pellets in 4 ml 100% IO. All prey resuspensions were at least 10^8^ cfu/ml. For the prey range tests, we plated the 10^-3^ to 10^-6^ dilutions of the filtrate. We incubated plates at 26°C and checked for plaques daily starting on day three until day seven. Plaque formation on any of these days was scored as positive for the prey range test. We repeated this procedure twice for each prey strain to obtain three biological replicates.

### Electron microscopy

To obtain BE01 samples for electron microscopy, we added a small amount of the −80°C stock of *Halobacteriovorax marinus* BE01 to 20 ml of 100% IO mixed with 1.5 ml of an *E. coli* #0057 overnight culture. After 48 hours of incubation at 26°C and 200 rpm, we placed formvar coated EM grids on 30 uL droplets of bacterial sample for 30 seconds to allow the bacteria to adhere to the formvar surface. We then transferred grids to 50 uL drops of 1% uranyl acetate in water for 1 minute. The grids were lifted from the drops of uranyl acetate and the excess stain wicked off with Whatman filter paper. The stained sample coated grids were air dried for 10 minutes and resulting specimens imaged with a Jeol CX 2000 transmission electron microscope.

### Sequencing and assembly of Halobacteriovorax marinus BE01 genome

To obtain genomic DNA for sequencing, we cultivated BE01 using *E. coli* #0057 as prey. We chose to use *E. coli* because there is extensive genome information available which would allow us to screen reads to remove prey bacteria reads if necessary. To make lysates for genomic DNA preparation, we added a small amount of the −80°C stock of BE01 to 20 ml of 100% IO mixed with 1.5 ml of an *E. coli* #0057 overnight culture. After two days of incubation, we examined the lysates for active predatory bacteria cells. We selected three lysates that appeared to have the highest ratio of predator to prey and pooled these. Because PacBio technology requires at least 10 μg of genomic DNA for library construction, we did not filter the lysates to avoid any potential loss of predatory bacteria cells. To extract genomic DNA, we used the Wizard Genomic DNA Purification Kit (Promega) with the protocol for Gram-positive and Gram-negative bacteria. We centrifuged the pooled lysates at 9100 rpm for 10 minutes and resuspended the pellet in 600 μl of Nuclei Lysis Solution. We then continued with the manufacturer’s instructions. At the final step, we left the genomic DNA at 4°C overnight. By Qubit 2.0, the genomic DNA was at 299 μg/ml.

Library construction and sequencing were performed at the Institute for Genome Sciences at the University of Maryland Baltimore on a Pacific Biosciences RSII instrument using P6-C4 chemistry. We launched an Amazon EC2 instance of SMRT Portal 2.3.0 to analyze and assemble the data. Two SMRT cells generated 93,922 post-filter polymerase reads (20,025 bp N50) and 151,636 subreads (10,161 bp N50). We performed *de novo* assembly using the RS_HGAP_Assembly.3 protocol [45] with default settings except for genome size, which we changed to 3.5 Mbp. This generated 93 contigs in the polished assembly. The largest contig was 3,413,657 bp and aligned to *Halobacteriovorax marinus* SJ by BLASTN. We used BLAST2Go [46] to align the other 92 contigs against the non-redundant database (restricted to Bacteria) with megablast to determine if any of the smaller contigs aligned to *Halobacteriovorax*.

To close the large *Halobacteriovorax* contig, we used Gepard [47] to identify overlaps between the ends of the contig, which indicated that the contig could be circularized. We used BLASTN alignments to specifically determine the overlap regions, which resulted in trimming 20,805 bp from the beginning of the contig. We then edited the trimmed contig so that the first nucleotide corresponded to the first nucleotide of the *dnaA* protein-coding sequence. To check the accuracy of the draft sequence at this stage, we aligned the PacBio reads against this draft sequence using the RS_Resequencing.1 protocol in SMRT Portal. The consensus sequence from this alignment had 542 differences compared to the draft sequence used as a reference.

To polish the sequence, we generated 150 bp paired-end Illumina reads. Library construction and sequencing (equivalent to 5% of a channel) were performed at IGS on an Illumina HiSeq. We filtered the resulting reads so that every base in each read pair was ≥Q25. This yielded 6,604,606 read pairs. We aligned these read pairs to the recalled draft sequence using bwa-mem [48], yielding 507x average coverage (with a minimum of 51x). We used samtools [49] to convert the alignment to a sorted and indexed bam file. Finally, we used Pilon [50] to identify corrections based on the Illumina data, which amounted to 372 small insertions. The corrected sequence generated by Pilon was deposited in GenBank as the complete chromosome of *Halobacteriovorax marinus* BE01.

### Genome annotation and analysis of gene content

We annotated the *H. marinus* BE01 genome initially using the NCBI Prokaryotic Genome Annotation Pipeline version 3.3. Because of the unusually high proportion of hypothetical proteins identified by PGAP (see Results), we submitted both BE01 and SJ chromosome sequences to RAST [18–20] in January 2017. We used classic RAST with the RAST gene caller and FIGfam Release70. We separately annotated RNA-coding genes using Infernal 1.1.2 [21]. Files of the RAST+Infernal annotations and the output from RAST bidirectional best hit analysis are available at the figshare repository (https://figshare.com/projects/Supporting_data_for_Halobacteriovorax_BE01_paper/24229). R code used to generate the synteny plot and the plot of amino acid identity for bidirectional best hits is available at the figshare repository (https://figshare.com/projects/Supporting_data_for_Halobacteriovorax_BE01_paper/24229).

### Modal codon usage analysis

To compare the modal codon usage frequencies, we used a freely available software package downloaded from http://www.life.illinois.edu/gary/programs/codon_usage.html [26].

## Additional files

### Additional File 1

Quicktime Movie .MOV

1000x phase-contrast microscopy of *Halobacteriovorax marinus* BE01 cells attacking *E. coli* ML35.

### Additional File 2

Excel spreadsheet

Classification of bacterial isolates used in prey range tests based on analysis of 16S rRNA gene sequences (>1000 bp) with three databases.

### Additional File 3

PDF

Neighbor-joining phylogeny of 16S rRNA gene sequences of four *Vibrio* isolates used in prey range tests, with *Enterovibrio norvegicus* (RDP accession LK391520) as outgroup. Distance matrix was constructed using dnadist in Phylip with Jukes-Cantor model of nucleotide substitution.

### Additional File 4

Excel spreadsheet

For individual genes discussed in the text, the RAST locus tag is matched with the corresponding PGAP locus tag to enable interested readers to quickly find relevant genes in the GenBank annotations.

## Declarations

### Ethics approval and consent to participate

Not applicable

### Consent for publication

Not applicable

### Availability of data and material

The genome sequence generated and analyzed during the current study is available as BioProject PRJNA343955, BioSample SAMN05806433 and GenBank accession CP017414. Other data and R code are available at the figshare repository (https://figshare.com/projects/Supporting_data_for_Halobacteriovorax_BE01_paper/242 29).

### Competing interests

The authors declare that they have no competing interests.

### Funding

This research was supported by an Institutional Development Award (IDeA) from the National Institute of General Medical Sciences of the National Institutes of Health under grant number P20GM103430 and funding from Providence College. Neither funder played a role in study design, data analysis or interpretation, or writing the manuscript.

### Authors’ contributions

BGE designed and performed experiments, analyzed data and wrote the manuscript. MKA designed and performed experiments. JAG performed negative staining and EM imaging. LEW designed experiments, analyzed data and wrote and edited the manuscript. All authors read and approved the final manuscript.

## Acknowledgments

We thank Nicole Cullen for isolating *Pseudomonas* #0042 and Sean O’Donnell for isolating *Acinetobacter* #0036. We thank Lisa Sadzewicz and Luke Tallon at the Institute for Genome Sciences at the University of Maryland Baltimore for sequencing services. We are grateful to Mark Martin for providing *E. coli* ML35, Brett Pellock for providing *E. coli* #0057 and Cameron Thrash for guidance on growth media. LEW thanks science Twitter for much useful advice and support.

## Authors’ information

BGE and MKA conducted this research as undergraduate researchers at Providence College.

